# Outsmarting refractory glioblastoma using *in silico* guided, phage display derived peptides targeting oncogenic EGFRvIII

**DOI:** 10.1101/2023.08.09.552576

**Authors:** Sunitha Kodengil Vettath, Giridharan Loghanathan Malarvizhi, Krishnakumar N Menon, Lakshmi Sumitra Vijayachandran

## Abstract

EGFRvIII mutation contributes to tumor aggressiveness and poor prognosis in multiple cancers. Despite being tumor-specific, EGFRvIII are not readily detected since they coexist with EGFR. Here, we developed novel, aqueous-dispersible peptides (1-15 kDa) that can detect and actively target EGFRvIII expressing refractory tumors, while sparing wild-type EGFR and normal astrocytes. Based on *in silico* molecular modeling, docking and biologics-based approaches, four EGFRvIII-targeting peptides were identified using phage display technology employing a) antibody fragments (AF), and b) random peptide library (RPL). Among RPL-based peptides, RB5 showed appreciable interactions with EGFRvIII receptor, as indicated by binding energy of -10.6 kcal/mol, *in silico. In vitro* binding studies using ELISA and immunocytochemistry further confirmed that RB5 could specifically target EGFRvIII+ glioblastoma cell, U87MG.ΔEGFR. Interestingly, AF-based peptide, H13, also showed considerable specificity towards U87MG.ΔEGFR. Importantly, these peptides exerted no cytotoxicity (MTS assay), or affected the downstream phosphorylation (western blot) in EGFRvIII cells. Thus, our study demonstrated that rationally designed peptide molecules can precisely target EGFRvIII tumors with minimal or no intracellular signaling interference. Owing to simple molecular architectures, these non-toxic, aqueous-dispersible peptides hold strong potential for conjugation with antitumor drugs and therapeutic nanoparticles for treating EGFRvIII positive refractory cancers.

## 1. INTRODUCTION

Epidermal growth factor receptor (EGFR) is a transmembrane glycoprotein that serves as a cell surface receptor. Under normal conditions, EGFR undergo dimerization and autophosphorylation upon ligand binding, and it regulates embryogenesis, cell survival, proliferation, differentiation, and migration [1,2]. However, when mutated, EGFR can lead to malignancies of the brain, breast, lung, ovary, pancreas, and colon [3–5]. Particularly, the deletion of exons 2-7 leads to generation of the constitutively active mutant receptor EGFRvIII, which contributes to extreme aggressiveness and poor prognosis in various cancers [6,7]. EGFRvIII mediates its downstream signaling primarily through PI3K-Akt pathway [8] leading to enhanced radio- and chemoresistance [9,10]. For instance, in 25-33% of glioblastoma patients, EGFRvIII mutation results in refractory tumor, thereby critically limiting the overall survival to 3-6 months, despite surgical resection, chemotherapy, and radiotherapy [11]. Thus, conventional therapies are often inadequate to address EGFRvIII mediated tumor recurrence. Further, targeted therapies using Chimeric Antigen Receptor T (CART) cells, vaccines, and antibodies provide only modest efficacy in EGFRvIII positive tumors, since the prognosis is limited by tumor recurrence, adverse effects, and insufficient tissue penetration, respectively [12,13]. Tyrosine kinase inhibitors targeting EGFRvIII also showed only modest efficacy owing to the activation of alternate signaling pathways and serious adverse effects [14]. Many anti-EGFRvIII monoclonal antibodies, such as DH1.1, DH8.3, 3C10, L8A4, and H10, show lack of target specificity and concerns of cross reactivity and hence not used in a therapeutic setting Importantly, due to their large size (M.W.: ∼160 kDa) and poor aqueous solubility, the utilization of monoclonal antibodies is limited by insufficient penetration and heterogeneous distribution in tumor tissue, leading to reduced efficacy [15,16]. In these contexts, peptide mediated active targeting appears promising for tissue-specific delivery of therapeutic cargos towards EGFRvIII positive tumors as their small size aids better penetration of conjugated drugs to the tumor interior. Further, peptides are better entities for functional group modifications and can be tailored to be hydrophilic, hydrophobic, or amphiphilic [17]. They are also relatively less immunogenic and possess better clearance *in vivo* due to their simple molecular architecture and low molecular weight (1-15 kDa), compared to antibodies that have high molecular weights [18].

The main objective of this work was to develop biocompatible, chemically stable, and aqueous-dispersible targeting peptides that can actively target EGFRvIII in drug-resistant, refractory tumors such as glioblastoma. To realize this, we employed *in silico* docking and phage display technology and identified novel peptides. These peptides were further validated for its EGFRvIII specificity using recombinantly expressed EGFRvIII ectodomain (ECD) and refractory tumor cells. The identified peptides had low molecular weight (1-15 kDa) and better aqueous dispersibility, compared to anti-EGFRvIII monoclonal antibodies. Further, the peptides had provisions for surface, site-specific, and functional group modifications to hydrophobically or covalently attach antitumor drugs, contrast agents, or therapeutic nanoparticles to specifically deliver therapeutic cargos towards EGFRvIII refractory tumors. The rationally developed peptides, even though actively targeted EGFRvIII, showed no significant toxicity towards normal astrocytes, or interfered with the downstream phosphorylation in EGFRvIII refractory glioblastoma cells, U87MG.ΔEGFR.

## 2. MATERIALS AND METHODS

We identified EGFRvIII binding peptides employing two approaches: (1) utilizing anti-EGFRvIII monoclonal antibody (mAb) based peptides sequences expressed on phage coat protein and (2) using a random peptide phage display library. In the first approach, peptides based on mAb 528 sequence were expressed on phage coat protein, which were thereafter screened against EGFRvIII expressing cells to identify EGFRvIII binding peptides. In the second approach, a commercially available random peptide library composed of 12-amino acid peptides displayed on M13 coat protein was screened tagainst to identify EGFRvIII binding peptides. The detailed information on the materials and methods used is provided in the supplementary text.

## 3. RESULTS

### 3.1. Identification, expression, and purification of mAb 528 based anti-EGFRvIII peptide

In order to identify EGFRvIII-targeting peptides, initially six putative peptide sequences located on the heavy and light chain regions of mAb 528 Fab (PDB ID: 2Z4Q) composed of one or more CDRs were selected. Thereafter, DNA sequences that encode these peptides were cloned into pCDisplay4 phagemid vector and peptide-displaying phages were generated. Fig. 1, A shows the binding of various peptide-displaying phages (H23, H13, H14, L12, L13, and L14) on glioma cells. Binding studies using whole cell phage ELISA revealed that H13 phage bound more specifically to EGFRvIII-positive glioma cell, U87-MG.ΔEGFR compared to EGFRvIII-negative cell, U87-MG. This binding was further substantiated by *in silico* docking studies (Supplementary Fig. 1) where H13 was found to interact strongly with EGFRvIII receptor (binding energy [B.E.]: -16.9 kcal/mol) than wild-type EGFR (-12.6 kcal/mol). Although H23, H14, and L12 phages also showed modest binding *in vitro* towards U87-MG.ΔEGFR compared to H13 phage, these phages showed appreciable binding towards U87-MG as well. Hence the specificity of binding towards the mutated cells (U87-MG.ΔEGFR) was significantly low for these phages.

**Figure 1.**
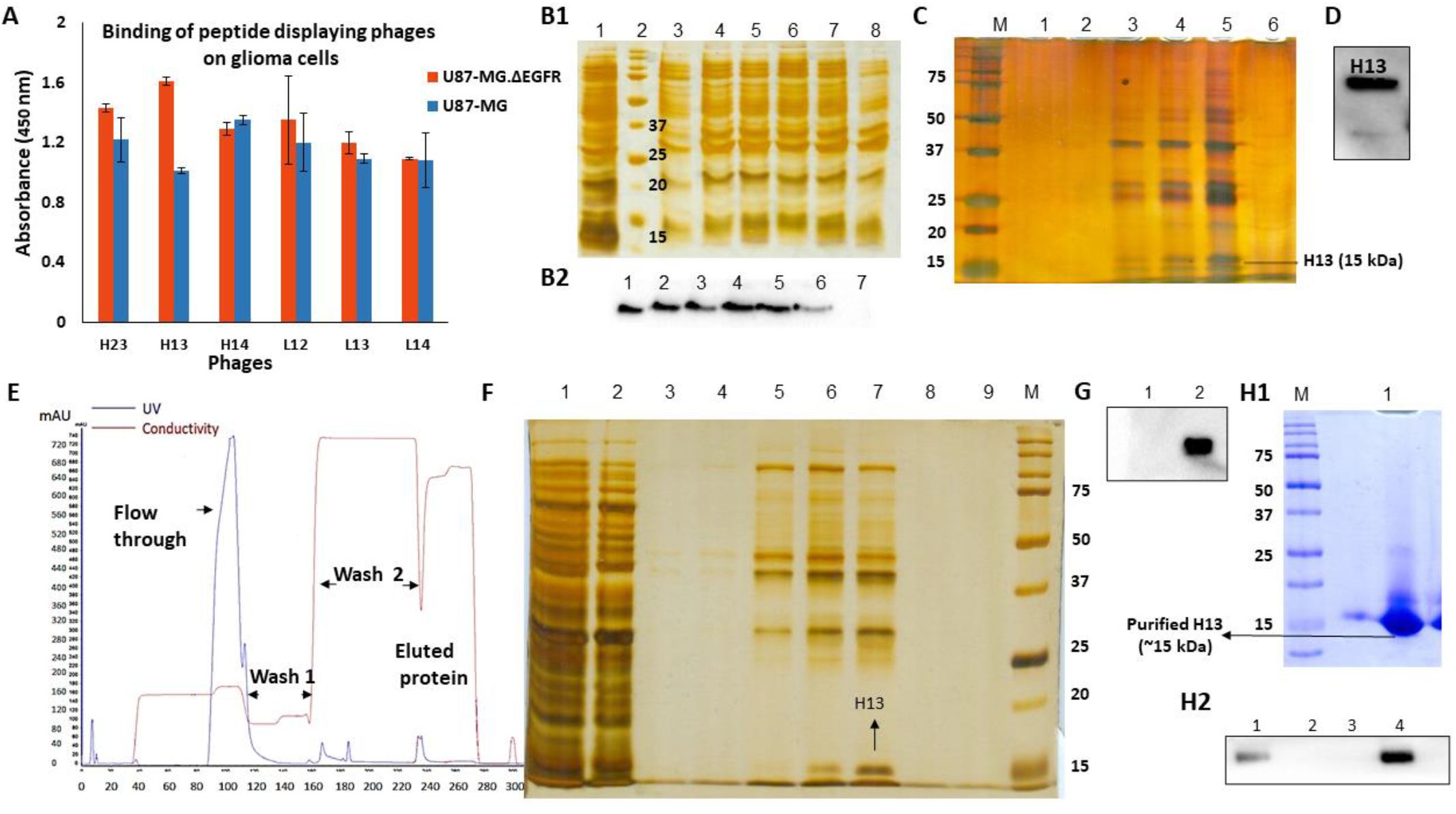
**A)** Whole cell phage ELISA of recombinant phages (displaying different fragments of mAb 528), on glioma cell lines. Values represent mean O.D@450nm ± S.D; **B1)** Total cell lysate of IPTG induced samples analyzed on 10% SDS PAGE-Lane 1: negative control (Top10F’ cells), Lane 2: Biorad Precision plus dual color protein ladder, Lane 3-8: pCDisplay4-H13 (Top10F’) induced with 1 mM IPTG at different temperatures and time points-Lane 3: 30°C, 3 hours, Lane 4: 30°C, 5 hours, Lane 5: 30°C, overnight, Lane 6: 37°C, 3 hours, Lane 7: 37°C, 5 hours, Lane 8: 37°C, overnight; **B2)** Western blot of total cell lysate using anti-His antibody (1:10000 dilution) shows the expression of H13. Lane 1: 30°C, 3 hours, Lane 2: 30°C, 5 hours, Lane 3: 30°C, overnight, Lane 4: 37°C, 3 hours, Lane 5: 37°C, 5 hours, Lane 6: 37°C, overnight, Lane 7: negative control (Top10F’ cells); **C)** Purification of anti-EGFRvIII peptide using Ni-NTA resin under hybrid conditions - Lane M: Biorad Precision plus dual color protein ladder, Lane 1: Wash 1, Lane 2: Wash 2, Lane 3: Elute 1, Lane 4: Elute 2, Lane 5: Elute 3, Lane 6: Elute 4. Elute 1, 2 and 3 were collected using elution buffer with 250 mM imidazole and elute 4 was collected using elution buffer with 500 mM imidazole; **D)** Western blot of elute using anti-His antibody: Elute 2; **E)** Affinity chromatography purification of anti-EGFRvIII peptide H13 from 0.25 L bacterial culture using AKTA START**; F)** 10% SDS PAGE analysis of AKTA purification-Samples were analyzed on gel. Lane 1: Lysate, Lane 2: Flow through, Lane 3: Wash, Lanes 4, 5, 6 and 7: Elutes 1-4 respectively, collected using elution buffer with 250 mM imidazole, Lanes 8 and 9: elutes collected using elution buffer with 500 mM imidazole; **G)** Western blot analysis of elutes collected with 250 mM imidazole, using Anti-his antibody. Lane 1: Elute 3, Lane 2: Elute 4; **H1)** SDS PAGE analysis of second step H13 purification. Samples were analyzed on 10% SDS PAGE gel. Lane M: Biorad Precision Plus dual color protein ladder, Lane 1: Purified H13; **H2)** Western blot analysis of elutes using Anti-his antibody. Lane 1: Elute from first round of purification, Lane 2: Flow through from second round purification, Lane 3: Wash, Lane 4: H13 from 2^nd^ round of purification.

### 3.2. Expression and purification of H13: Ni affinity chromatography

In the next step, for studying the binding of isolated H13 peptide in EGFRvIII-positive cells, purification was performed using Ni affinity chromatography. For this, at first pCDisplay4-H13 phagemid was transformed into *E. coli* Top10F’ cells and the expression of H13 peptide was studied. Fig. 1, B1 shows the SDS PAGE of total cell lysates of IPTG induced H13-Top10F’ cultures at different temperatures (30°C and 37°C) and incubation times (3 h, 5 h, and 12 h). Lane 1 shows the total cell lysate of Top10F’ cells (negative control) and lanes 3-8 show the IPTG induced cell lysates. Although feeble-to-strong bands were observed in all IPTG induced cell lysates at 15 kDa (H13 peptide), negative control also showed a relatively strong band at the same molecular weight. Therefore, western blot was performed using anti-Histidine antibody that revealed the expression of H13 in all the IPTG induced samples (Fig. 1, B2) with no band in the negative control, which clearly confirmed H13 peptide expression (Fig. 1, B2, lane 7). Notably, the expression of H13 was better when induced at 37°C for 3 h and 5 h (lane 4, 5, respectively). Subsequently, H13 was purified from 50 ml culture using Ni-NTA affinity chromatography. Fig. 1, C shows the wash and elution fractions of H13 analyzed on SDS PAGE gel. Elution fractions in lanes 3, 4, 5, collected using elution buffer containing 250 mM imidazole showed the presence of H13 peptide, which was further confirmed using western blot (Fig. 1, D). Fig. 1, E shows the chromatogram of the purification of H13 from 250 ml culture using AKTA START FPLC system, and Fig. 1, F shows the SDS PAGE profile of different purification steps, where the lanes 6 and 7 shows the presence of H13 peptide collected using 250 mM imidazole containing elution buffer. The presence of H13 was confirmed using western blot (Fig. 1, G). Fig. 1, H1 shows the SDS PAGE of purified H13 peptide after second step of purification whose identity was further confirmed using western blot (Fig. 1, H2).

### 3.3. Binding of H13: Immunocytochemistry and ELISA

Following the purification of H13, binding of the purified H13 peptide was studied in glioma cells (U87-MG.ΔEGFR, U87-MG) and normal astrocytes (CTX TNA2), which do not express EGFRvIII, using immunocytochemistry. Fig. 2, A1, B1, and C1 shows the binding of H13 peptide in U87-MG.ΔEGFR, U87-MG, and CTX TNA2, respectively, and their enlarged images are shown in Fig. 2, A2, B2, and C2. An apparent binding of H13 peptide was evident in EGFRvIII-positive cell, U87-MG.ΔEGFR, as indicated by the brown staining in these cells (Fig. 2, A1 and A2). In contrast, EGFRvIII-negative cell, U87-MG, and normal astrocytes, CTX TNA2, did not show the brown staining pattern after incubating with H13 peptide. For further confirming the binding of H13 peptide towards EGFRvIII, we have quantitatively studied the binding in recombinant EGFRvIII ECD and glioma cells. Fig. 2, D shows ELISA results of binding of H13 on recombinant ECD. As can be noticed, compared to the negative control (BSA), there was a concentration dependent increase in the binding intensity of H13 in EGFRvIII ECD. We further studied the binding of H13 in glioma cells and compared them with that of mAb 528, with appropriate controls. We have used three different cells for this study: C6 glioma (EGFR- and EGFRvIII-negative); U87-MG (EGFR-positive but EGFRvIII-negative) and U87-MG.ΔEGFR (EGFR- and EGFRvIII-positive cell). As can be seen in Fig. 2, E and F, both H13 peptide and mAb 528 did not show an appreciable binding towards C6 glioma cells, since these cells lack EGFR as well as EGFRvIII receptors. H13 peptide showed a modest binding with U87-MG cells, which was not statistically significant (Fig. 2F). However, when H13 peptide was treated in U87-MG.ΔEGFR cells, compared to mAb 528, a strong and intensely specific binding was clearly evident in these EGFRvIII-positive cells.

**Figure 2.**
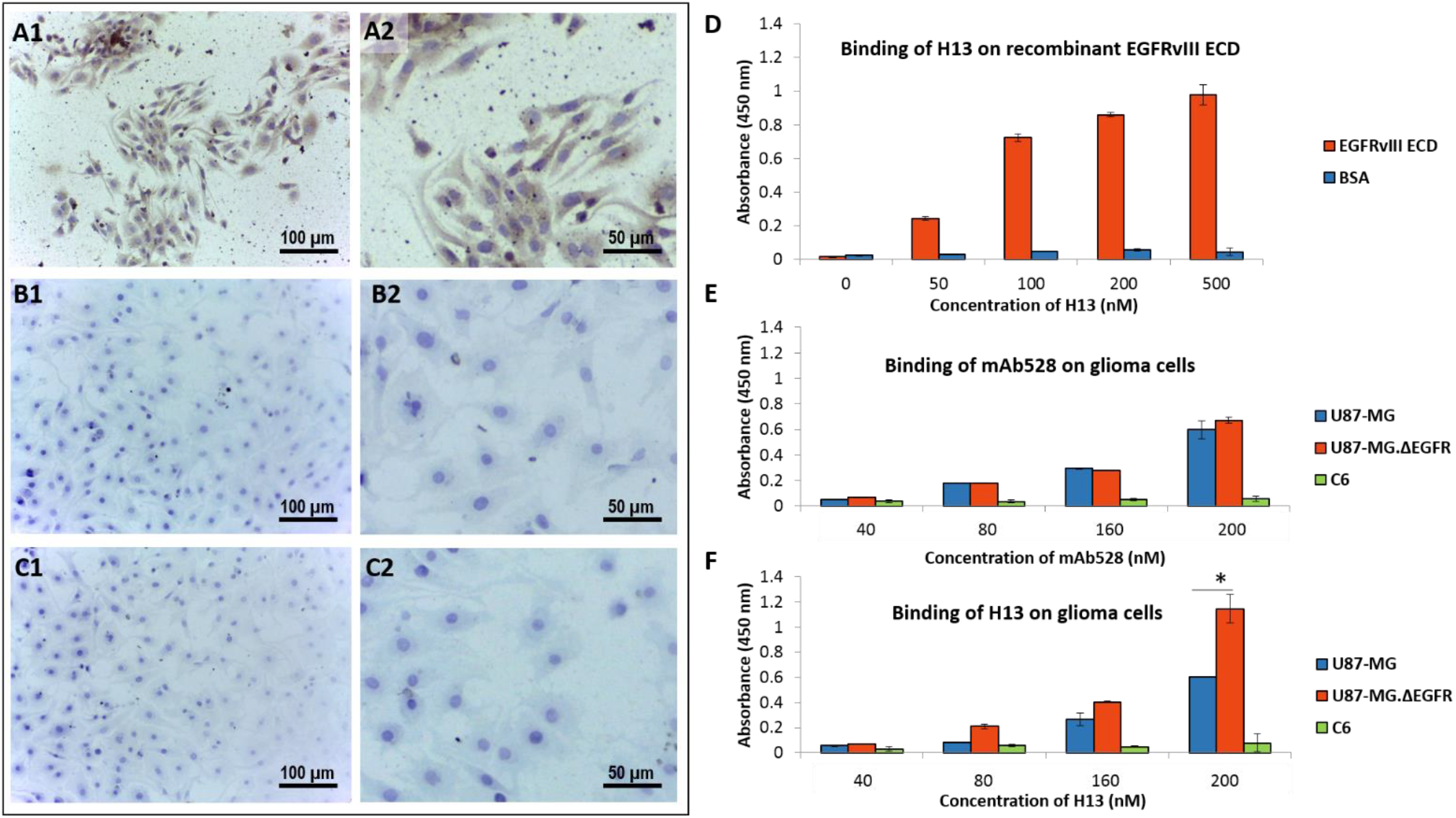
Immunocytochemical staining of EGFRvIII expressing cells with purified H13 showing binding of H13 to U87-MG.ΔEGFR (by DAB, 200×), detected using anti-Histidine antibody: **A1)** U87-MG.ΔEGFR; **B1)** U87-MG; **C1)** normal astrocyte cell line CTX–TNA2. The absence of staining in U87-MG and normal astrocyte cell line CTX– TNA2 can be clearly noticed. Fig. **A2, B2** and **C2** represents the zoomed versions of Fig. A1,B1, and C1 respectively; **D)** ELISA of H13 binding on recombinant EGFRvIII ectodomain. Graph represents O.D at 450 nm±S.D; **E)** Whole cell ELISA of mAb 528 and (F) purified H13 binding on cell lines. Graph represents O.D at 450 nm±S.D.). (*) P<0.05 significant.

### 3.4. Cell viability and phosphorylation studies: H13

In the next step, we investigated whether H13 peptide has potential to affect cell viability or inhibit phosphorylation of EGFRvIII and its downstream signaling kinase, PI3K. MTS assay revealed that H13 peptide did not cause a significant reduction in the viability or proliferation of U87-MG.ΔEGFR cells or normal astrocytes, CTX TNA2 (Fig. 3, A). Although H13 peptide could recognize and bind to these mutant receptors, western blot results revealed that the phosphorylation of EGFRvIII (Fig. 3, C) and PI3K (Fig. 3, D) was not significantly different after treatment with H13. Fig. 3, B shows the representative western blot images of EGFRvIII and PI3K phosphorylation in U87-MG.ΔEGFR cells after treating with H13 or mAb 528 (control).

**Figure 3.**
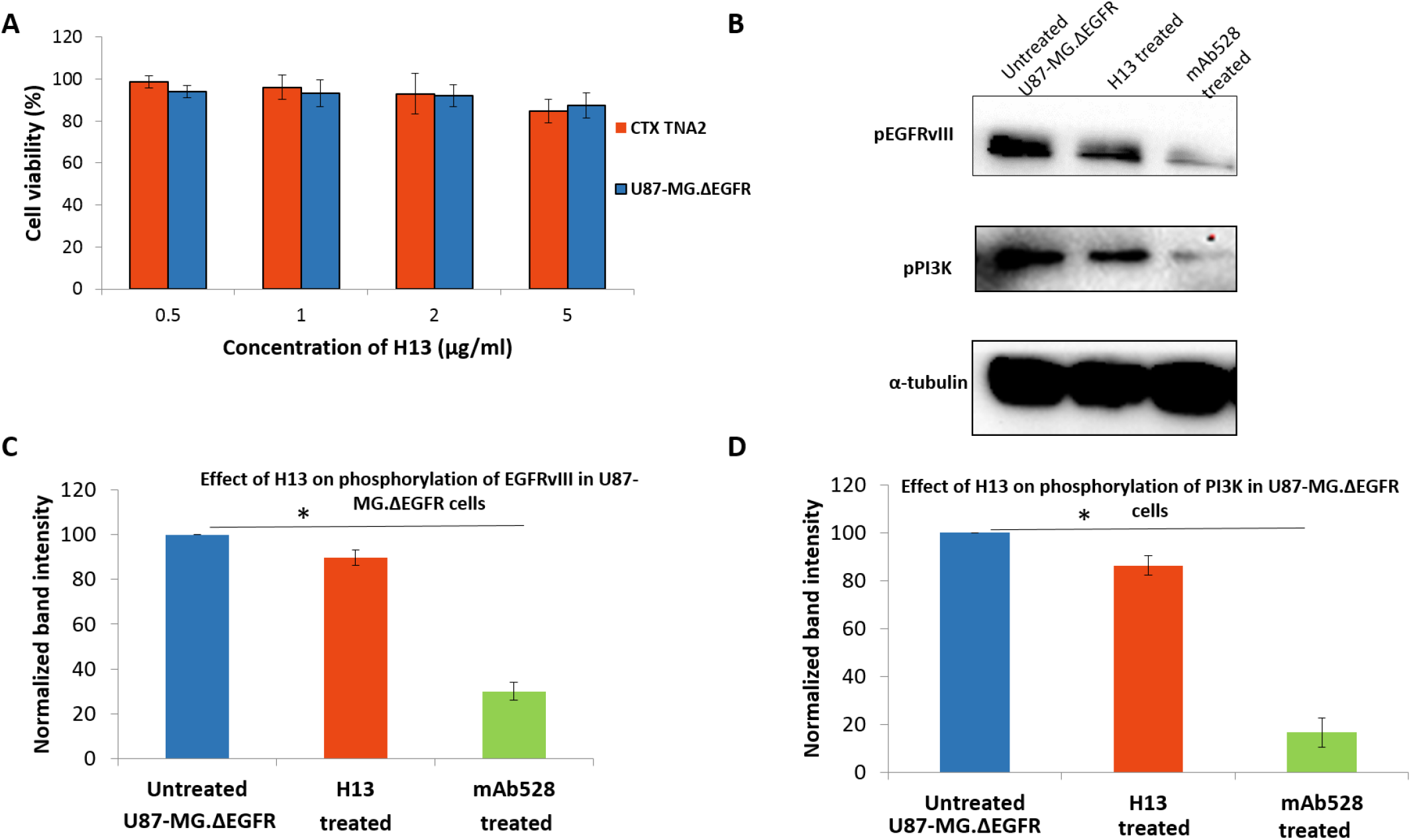
**A)** Antiproliferative effect of purified H13 peptide on cell lines. Bars represent percentage of cell viability [Mean (O.D of treated cells/Mean O.D of cell control) ×100] ± S.D (n=3); **B)** Effect of H13 and mAb 528 on the phosphorylation of EGFRvIII and PI3K in U87-MG.ΔEGFR cells - representative western blot image; **C)** Effect of H13 and mAb 528 on the phosphorylation of EGFRvIII; **D)** Effect of H13 on the phosphorylation of PI3K

### 3.5. Identification of EGFRvIII binding peptides: RPL-based screening

In order to identify alternate peptides that interact with EGFRvIII with more specificity and affinity, we utilized phage displayed random peptide library wherein many peptides are displayed on phage and provides the opportunity to identify specific and high affinity peptides shorter (<2 kDa) than H13 (∼15 kDa). Short peptides exhibit better tumor penetration potential and chemical synthesizability, which are beneficial for clinical translation. To realize this, biopanning of Ph.D.^TM^-12 phage display peptide library was executed for obtaining EGFRvIII-specific lead phages. Table 1 shows the input-output ratios that denote an obvious enrichment of EGFRvIII-binding peptides after three rounds of biopanning in subtractive and reverse biopanning methods. After the second round, there was a 5-fold and 100-fold increment in the phage output, which after the third round increased substantially resulting in 80 and 150-fold increment in subtractive and reverse biopanning, respectively. Next, whole cell ELISA of phages, amplified from the acid (cell surface bound) and output fraction (cell interior) of subtractive and output fraction from reverse biopanning, on U87-MG.ΔEGFR cells was performed. For the selection of EGFRvIII-specific phages, an arbitrary value of 1 was assigned as the cut-off point for O.D450nm. In the case of subtractive biopanning, for the acid fraction, no phages crossed the cut-off point; however, for the output fraction, three phages (SB6, SB12, and SB31) showed O.D values greater than 1 (Fig. 4, A). In the case of reverse biopanning, for the output fraction, two phages (RB5 and RB8) showed O.D values greater than 1 (Fig. 4, B). The single stranded DNA of selected EGFRvIII-specific phages were isolated, visually verified using 1% agarose gel (Fig. 4, C) and were submitted for sequencing. Sequence of three peptides SB12, SB31, and RB5 were obtained.

**Figure 4.**
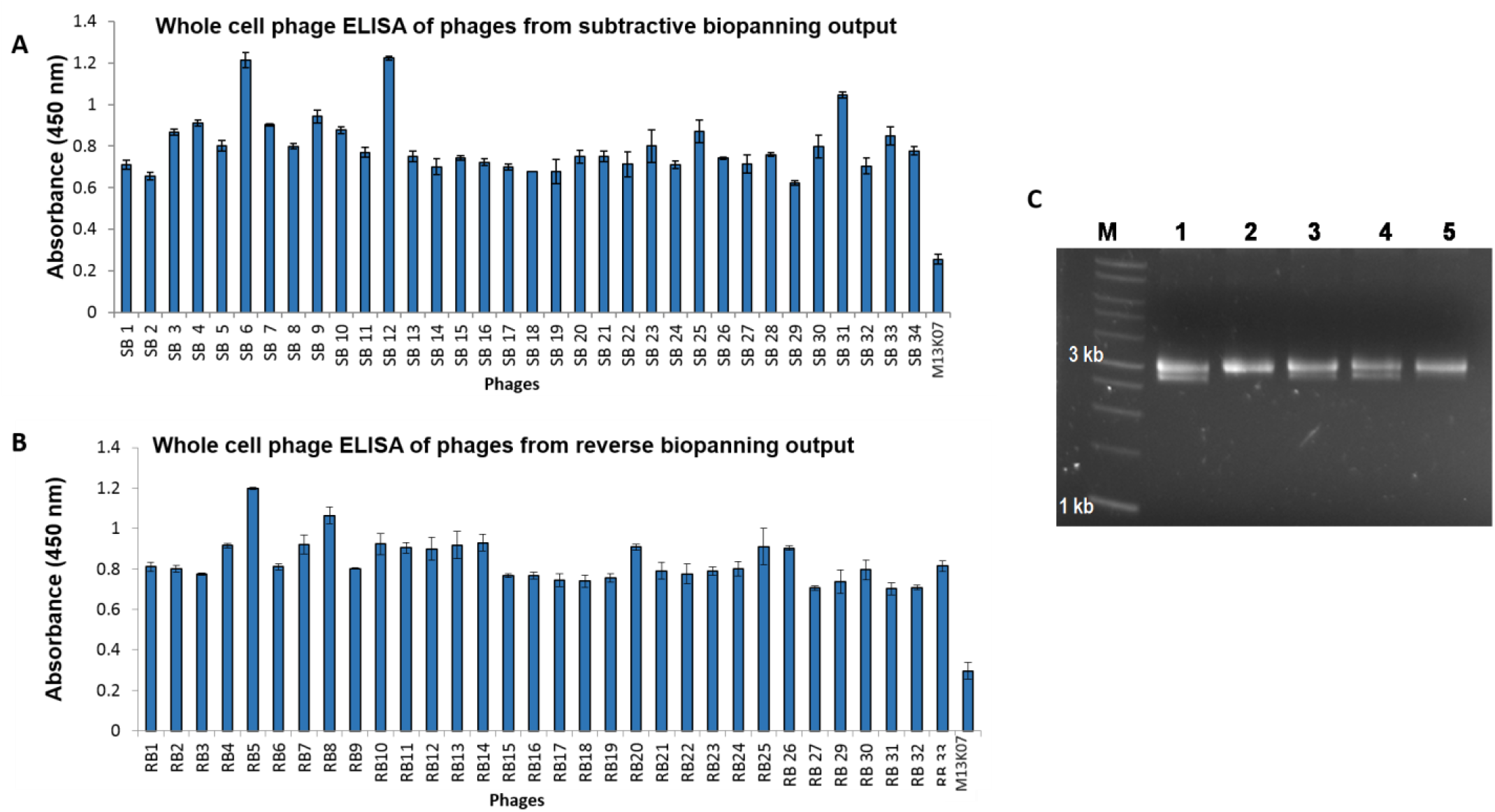
**A)** Whole cell ELISA of phages amplified from subtractive biopanning output. Values represent mean O.D at 450nm ± SD (n=3); **B)** Whole cell ELISA of phages amplified from reverse biopanning output. Values represent mean O.D at 450nm ± SD (n=3). M13K07 helper phage (MK) was used as the negative control; **C)** Agarose gel electrophoresis of ssDNA isolated from selected phages. Lane M: 1kb plus ladder; Lane 1-5: ssDNA from SB6, SB12, SB31, RB5, and RB8 phages respectively; **D)** Sequence of peptides displayed by the selected phages, derived from ssDNA sequencing results.

**Table 1:**
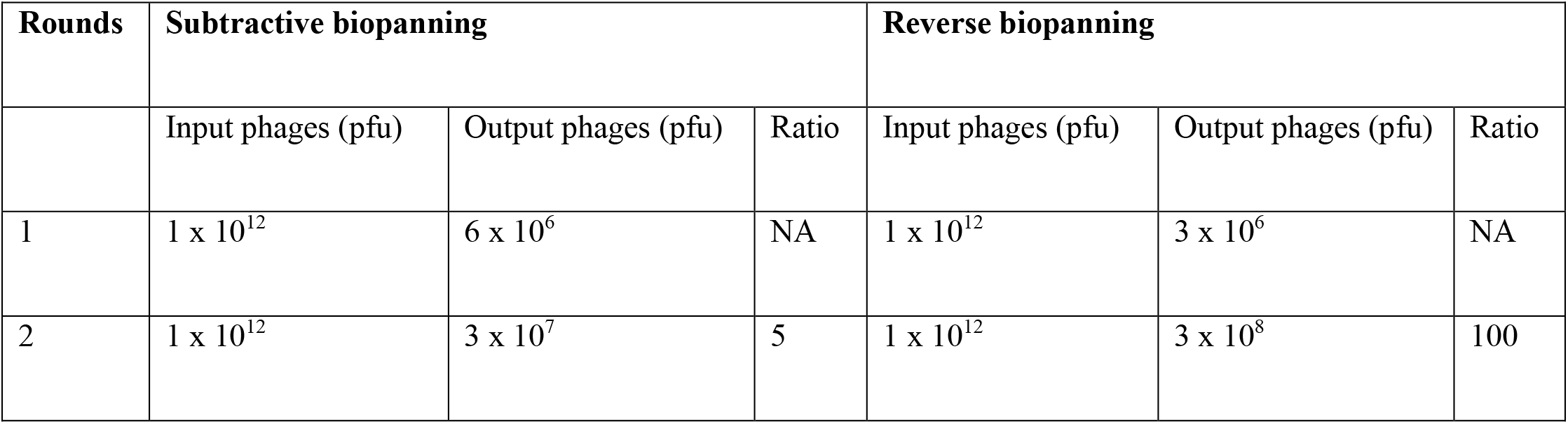
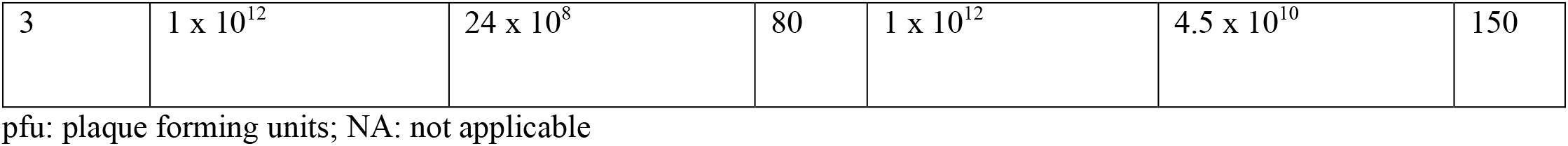
Ratio of output after each biopanning round in subtractive and reverse biopanning.

### 3.6. *In silico* molecular modeling and docking of EGFRvIII and peptides

Since one of our main objectives was to study the active targeting capability of peptides towards EGFRvIII, prior to *in vitro* realization, *in silico* molecular modeling and docking simulations were employed to get the preliminary information on whether the identified peptides can indeed bind this mutated receptor. For this, EGFRvIII ECD was homology modeled first, since its crystal structure is not yet experimentally resolved. Thereafter, 3D structure of the peptides were separately modeled and docked with EGFRvIII ECD to study the targeting capability. Fig. 5, A shows the crystal structure of unmutated or wild-type EGFR (PDB ID: 1IVO), the parent receptor of EGFRvIII. The residues labeled in red denotes the deleted regions of the wild-type receptor, and the undeleted or retained residues that constitute the EGFRvIII mutant is displayed in green. Fig. 5, B shows the homology modeled structure of EGFRvIII ECD that results from in-frame deletion of 801 base pairs spanning exons 2-7 of the coding sequence of the wild-type EGFR. This deletion removes 267 amino acids from the extracellular domain, while creating a junction site between exons 1 and 8 and a new glycine residue. The EGFRvIII ECD model obtained after side chain refinement and energy minimization was validated using Ramachandran plot (Fig. 5, D). Thereafter, the various ligand-binding pocket residues of EGFRvIII ECD were identified. Interestingly, we found that at least 8 potential ligand-binding pockets are present in the ECD region of EGFRvIII (labeled in distinct colors for better visualization) (Fig. 5, C). Further functional annotation indicated that Pro 41, Ile 289, Cys 291, Lys 36, and Cys 38 in EGFRvIII ECD are involved in metal ion-binding activity (Supplementary Table 2). Subsequently, the tertiary structures of the 3 peptides (SB12, SB31, and RB5) were modeled *in silico* (Fig. 5, E, F, G). It was found that the peptides SB12 and SB31 formed right-handed alpha helices with terminal loops. In contrast, RB5 model displayed a loop structure. The structure of the 3 modeled peptides were also validated using Ramachandran plot, and all the models were found to lie well within the allowed regions (Supplementary Fig. 2). Thereafter, SB12, SB31, and RB5 peptides were individually docked with EGFRvIII ECD (Fig. 5, I, J, K). The results revealed stable binding of all the three peptides with the mutant receptor, as indicated by their appreciable binding energies (Fig. 5, H). However, interestingly, among the 3 peptides, RB5 peptide showed a relatively stable binding towards EGFRvIII ECD as indicated by its lowest binding energy (B.E.: -10.6 kcal/mol). In the next step, the nature of chemical interactions between the RB5 peptide and the EGFRvIII mutant receptor was studied. Fig. 5, L shows the closely interacting amino acid residues of EGFRvIII ECD with RB5 peptide. The complete list of closely interacting residues of the 3 peptides with EGFRvIII ECD is provided in the supplementary information (Supplementary Fig. 3).

**Figure 5.**
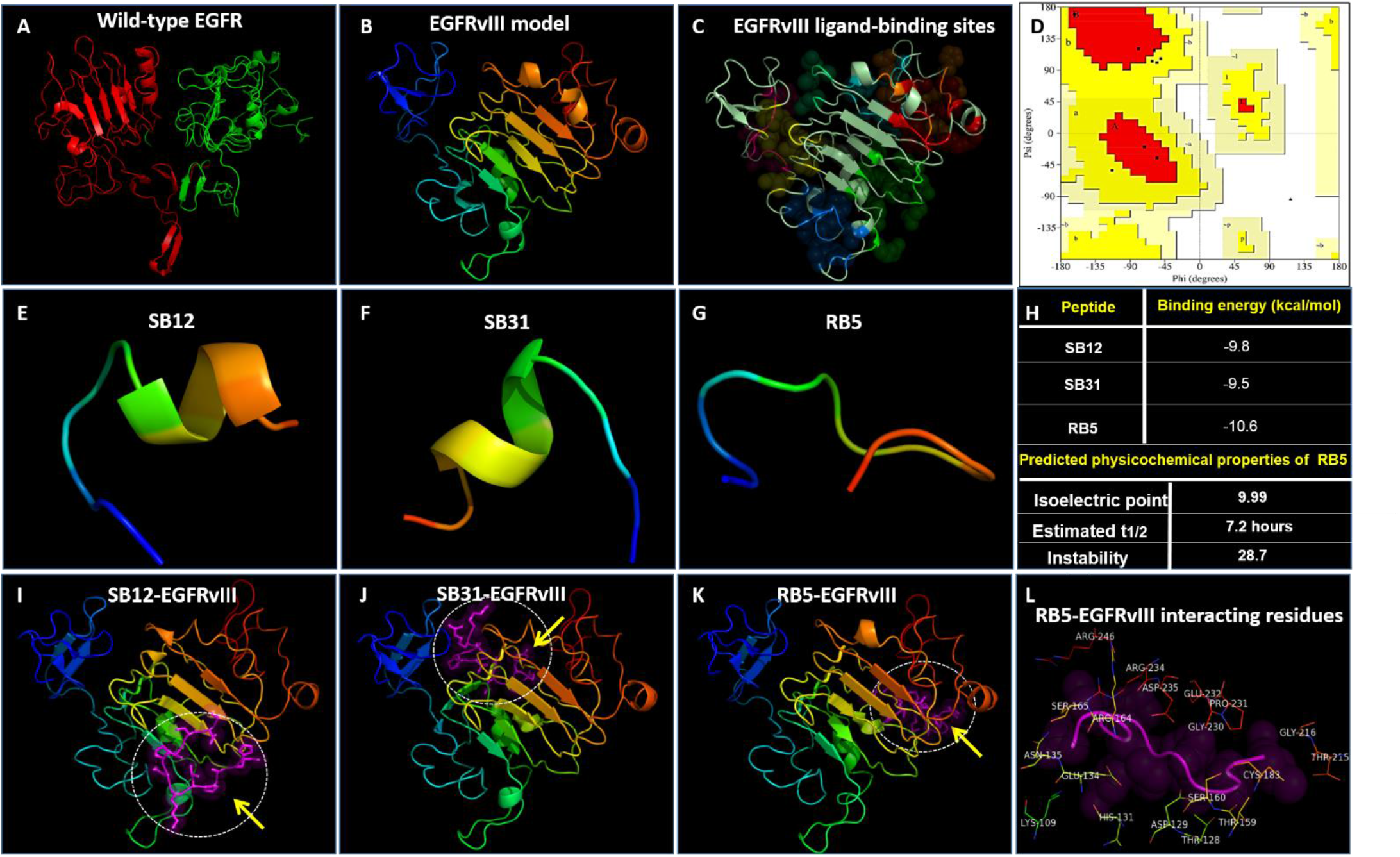
**A)** Crystal structure of wild-type EGFR shows the unaltered (red) and mutated regions (green) of the receptor; **B)** Homology model of EGFRvIII receptor; **C)** EGFRvIII shows potential ligand-binding sites (displayed in distinctly colored spheres); **D)** Structural verification of EGFRvIII using Ramachandran plot; Modeled structures of EGFRvIII-targeting peptides **E)** SB12, **F**) SB31, and **G)** RB5; **H**) *In silico* docking associated binding energies of EGFRvIII-targeting peptides with EGFRvIII, and estimated physicochemical characterization of RB5 peptide; *In silico* docking simulations of **I)** SB12, **J)** SB31, and **K)** RB5 peptides with EGFRvIII receptor; **L)** Closely interacting amino acid residues of RB5 peptide with EGFRvIII receptor in RB5-EGFRvIII docked complex (< 3 Å distance).

### 3.7. Cell viability and phosphorylation studies: SB12, SB31, RB5

Following the *in silico* studies, SB12, SB31, and RB5 peptides were synthesized at Genscript (as mentioned in the methods section) and studied the cytotoxicity of all the three peptides in glioma cells. Interestingly, MTS assay revealed no significant cell death for any of the tested peptides (Fig. 6, A), and they did not interfere with the phosphorylation of EGFRvIII and its downstream kinase, PI3K (Fig. 6, C and D). Fig. 6, B shows the representative western blot images of EGFRvIII and PI3K phosphorylation in U87-MG.ΔEGFR cells after treating with these peptides.

**Figure 6.**
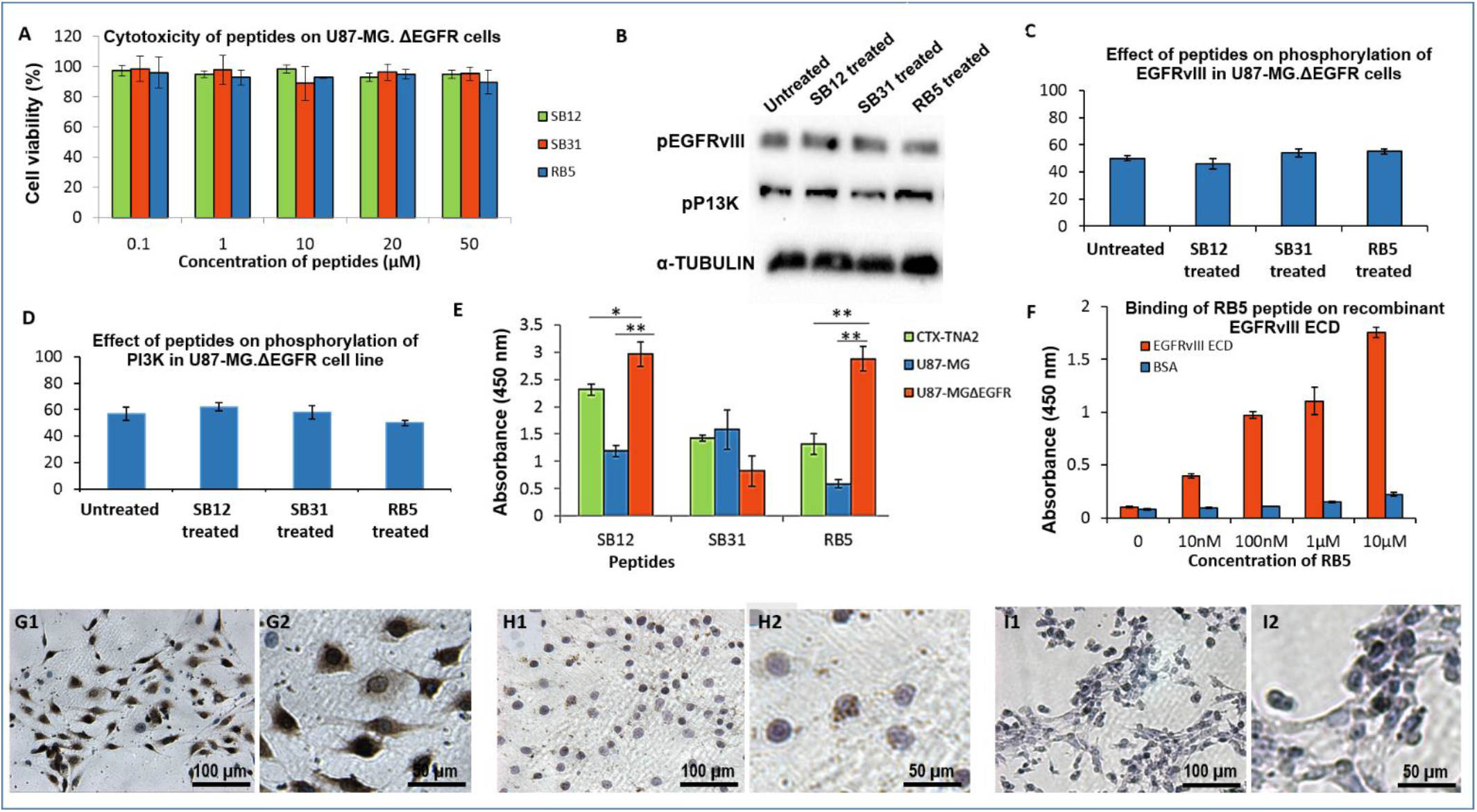
**A)** Cytotoxicity of peptides SB12, SB31, and RB5 on U87-MG.ΔEGFR cells. Values represent [average (O.D -average O.D_blank_)/average O.D_cell control_ -average O.D_blank_] X 100 ±S.D; **B)** Representative western blot for mechanistic studies of peptides; **C)** Effect of peptides on the phosphorylation of EGFRvIII; **D)** Effect of peptides on the phosphorylation of PI3K; **E)** ELISA of biotinylated peptide binding on cell lines. Values represent average [O.D at 450nm-average O.D of blank] ± S.D. Student’s t-test was used to test statistical significance of RB5 and SB12 peptide binding in various cell lines; (U87-MG.ΔEGFR versus CTX-TNA2, U87-MG.ΔEGFR versus U87-MG). (*) P<0.05 significant, (**) P<0.01 significant; **F)** ELISA of RB5 binding on recombinant EGFRvIII ectodomain. Graph represents O.D at 450nm±S.D; **G1-I2**) Immunocytochemical staining of RB5 peptide binding on cell lines (by DAB). **G1)** U87-MG.ΔEGFR; **H1**) U87-MG; **I1)** normal astrocyte cell line CTX–TNA2. Fig **G2**, **H2** and **I2** represent the zoomed versions of Fig G1, H1, and I1 respectively.

### 3.8. Binding of SB12, SB31, and RB5 on glioma cells: ELISA

Following that, we studied the binding of SB12, SB31, and RB5 peptides using ELISA in U87-MG.ΔEGFR, U87-MG, and CTX TNA2 cells (Fig. 6, E). The results revealed better binding of both SB12 and RB5 in the refractory glioma cells, U87-MG.ΔEGFR, compared to SB31. With regard to the specificity of EGFRvIII binding, although both RB5 and SB12 showed comparable binding towards U87-MG.ΔEGFR cells, RB5 showed lesser non-specific binding towards U87-MG and CTX TNA2 cells.

### 3.9. Binding of RB5 on EGFRvIII ECD and EGFRvIII-positive glioma cells

Based on the above observation, we further confirmed the binding quantitatively in recombinant EGFRvIII ECD using ELISA (Fig. 6, F). As can be noticed, compared to the negative control (BSA), there was a concentration dependent increase in the binding intensity of RB5 in EGFRvIII ECD. Following that, binding of RB5 peptide was studied in glioma cells (U87-MG.ΔEGFR, U87-MG) and normal astrocytes (CTX TNA2) using immunocytochemistry. Fig. 6, G1, H1, and I1 shows the binding of RB5 peptide in U87-MG.ΔEGFR, U87-MG, and CTX TNA2, respectively, and their enlarged images are shown in Fig. 6, G2, H2, and I2. An apparent binding of RB5 peptide was evident in EGFRvIII-positive cell, U87-MG.ΔEGFR, as indicated by the intense brown staining in these cells (Fig. 6, G1 and G2). However, EGFRvIII-negative cell, U87-MG, and normal astrocytes, CTX TNA2, did not show any staining after incubating with RB5 peptide. This revealed that the synthesized novel RB5 peptide has increased specificity and can be used for targeting EGFRvIII-positive cells without affecting the EGFRvIII-negative cells or normal astrocytes.

## 4. DISCUSSION

EGFRvIII mutations that are common in glioblastoma-Grade IV confer tumor refractivity and intrinsic drug resistance. EGFRvIII have been detected in solid tumors of the colon (34%), esophagus, ovary (75%), breast (78%), head and neck (42%), and non-small cell lung carcinomas [19–22], and the overexpression of EGFRvIII in high-grade glioblastoma multiforme has been consistently confirmed [23].

In this study, we developed EGFRvIII-specific peptides that can actively target EGFRvIII in drug-resistant tumors such as glioblastoma. Further, the possibility of developing non-toxic and aqueous-dispersible targeting peptides was also explored. For this, peptides that can specifically target EGFRvIII were identified using two different technologies: a) antibody fragment based, b) random peptide phage display library. *In silico* molecular modeling and docking simulations were employed to rationally identify candidate peptides that can specifically and stably target EGFRvIII receptors. The nature of chemical interactions (hydrophobic, hydrogen bonds, electrostatic) between the peptide and the mutant receptor were quantitatively studied.

At first, we have studied the binding of the six peptide-displaying phages (H23, H13, H14, L12, L13, and L14) on glioma cells using whole cell phage ELISA. The results revealed that H13 phage can bind more specifically to EGFRvIII-positive glioma cells than to EGFRvIII-negative cells, which was further substantiated by the *in silico* H13-EGFRvIII docking analysis. We found the expression of H13 was high at 37°C when incubated for 3 h and hence followed these conditions for purification. We purified the protein using Ni-NTA chromatography under optimized hybrid conditions that involved cell lysis, binding, and washing, initially under denatured conditions, followed by a final washing and elution performed in native conditions. This hybrid approach was an important strategy that practically enabled the better enrichment of the peptide and binding on EGFRvIII-positive cells. Thus, H13-displaying phages were identified and H13 peptide was, expressed, purified and finally verified using western blot. Immunocytochemistry experiments conducted using H13 indicated that it can specifically target EGFRvIII-positive cells without affecting EGFRvIII-negative cells or normal astrocytes. ELISA results revealed concentration-dependent increase in the binding intensity of H13 on recombinant ECD [24]. Further *in vitro* binding studies in cells using ELISA revealed that, compared to mAb 528 (control), recombinant H13 peptide has more specificity in binding to EGFRvIII, as indicated by the increase in the binding intensity in U87-MG.ΔEGFR cells. Studies on the cell viability and phosphorylation of EGFRvIII and its downstream kinase signaling molecule, PI3K, revealed no significant reduction in the viability of U87-MG.ΔEGFR cells or normal astrocytes after treating with H13 peptide; neither the phosphorylation was affected.

We further attempted to identify alternative short peptides from phage display libraries as they may have a better tumor tissue penetration *in vivo*. Five EGFRvIII-specific phages were identified from the Ph.D.^TM^-12 phage display peptide library after three rounds of biopanning and whole cell ELISA. The sequence of the displayed peptide in three phages were identified (SB12, SB31, and RB5); the other two phages SB6 and RB8 showed no inserts. We believe that SB6 and RB8 phages had lost their peptide owing to their instability, consequently deleted during repeated amplification. Sequence analyses using BLASTP confirmed that the EGFRvIII-binding peptides SB12, SB31, and RB5 are novel, as revealed by the absence of sequence identity with any other peptides in the NCBI non-redundant protein sequence database.

Prior to studying the binding of SB12, SB31, and RB5 peptides with EGFRvIII ECD *in vitro*, *in silico* molecular modeling and docking simulations were performed. The simulation results indicated that EGFRvIII ECD has at least 8 ligand-binding pockets having potential to bind SB12, SB31, or RB5 peptides. This was later confirmed by the docking experiments where although all the 3 peptides bound with EGFRvIII ECD, RB5 peptide showed the most energetically-favorable binding with EGFRvIII. Further investigations on the nature of chemical interactions between the closely interacting amino acid residues of RB5 and EGFRvIII revealed that the peptide-mutant receptor interactions are stabilized primarily by the combination of hydrogen bonds, electrostatic interactions, and to an extent by hydrophobic and Van Der Waal’s forces (Supplementary Fig. 3). Further functional annotation of modeled EGFRvIII ECD indicated that the mutant receptor has potential to bind with ionic compounds, particularly, metallic cations and transition metals such as iron, copper, nickel, and zinc, thereby possibly modulating metal ion-dependent cellular metabolic processes through downstream signal transduction (Supplementary Table 2). Possibly, this cationic metal binding property is one reason behind the pathogenesis and proliferation of heavy metal induced-cancers that express EGFR/EGFRvIII, as reported elsewhere [25]. However, the metal binding property of EGFRvIII ECD might be a double-edged sword. On one hand, it can contribute to pathogenesis by activating metal-dependent oncogenic kinases, proteins or enzymes, while on the other hand, it can provide potential opportunity for using metal-based antitumor targeting and therapy against EGFRvIII associated tumors [26,27]. Nevertheless, this is the first report that gives an indicative information on metal binding by EGFRvIII ECD from the context of tumorigenesis, as well as for aiming a possible antitumor therapy using metals for combating EGFRvIII-positive cancers. Similarly, our studies also indicated that EGFRvIII ECD indeed has potential for appreciable drug/ligand binding, despite its ligand-independent constitutive phosphorylation activity. Nevertheless, whether targeting these ligand-binding pockets can have any therapeutic value needs to be investigated.

Cytotoxicity studies revealed no significant cell death for SB12, SB31 or RB5 peptides in glioma cells. Further, western blot revealed that the peptides did not affect the phosphorylation of EGFRvIII or PI3K. ELISA-based binding studies conducted in U87-MG.ΔEGFR cells revealed appreciable binding for both SB12 and RB5. However, RB5 peptide showed statistically significant EGFRvIII-specific binding. This result was consistent with the *in silico* docking wherein RB5 peptide displayed the most stable binding towards EGFRvIII ECD. Interestingly, RB5 showed relatively less non-specific binding towards wild-type glioma cells and normal astrocytes. Similarly, ELISA-based binding studies conducted in purified recombinant EGFRvIII ECD and immunocytochemistry results revealed EGFRvIII-specific binding of RB5.

Owing to the clinical success of peptides, targeting-peptides have been approved for the diagnosis and treatment for several cancers. For instance, Degarelix, a hormone-modulating synthetic therapeutic peptide that binds to gonadotropin-releasing hormone (GnRH) receptors, appears to be promising for treating advanced prostate cancer. Similarly, radioactive Lutetium (Lu 177) conjugated with somatostatin receptor-targeting peptide, dotatate, is used in clinics for treating gastroenteropancreatic neuroendocrine tumors [28]. Thus, peptides are excellent candidates for both targeting and therapy for difficult-to-treat tumors.

Our findings are significant as the peptides have the potential to selectively deliver drug payloads and eliminate EGFRvIII positive refractory cancer cells. In addition to their ease of synthesis, these peptides could also serve as valuable tools for identifying EGFRvIII-expressing cancers, highlighting their versatility and potential in precision cancer theranostics. For instance, superparamagnetic iron oxide nanoparticles (SPIONs) loaded EGFRvIII-targeting peptides have been used as diagnostic tools for dual-modality imaging of intracranial EGFR-positive glioblastoma [29]. Similarly, EGFR/EGFRvIII dual-targeting peptide-mediated system has been successfully used as drug delivery vehicles for enhancing the therapeutic efficacy in glioma [30]. In this study, employing phage display technology, we have successfully developed two novel water-soluble peptides, H13 and RB5, those can precisely and specifically target EGFRvIII receptors in refractory glioma cells while having relatively less non-specific uptake towards wild-type EGFR-positive cells and normal astrocytes. H13 peptide identified based on antibody although specifically targeted EGFRvIII, has relatively large size (∼15 kDa) compared to RB5 peptide (∼1.2 kDa). From practical view point, this small size provides provisions for facile synthesis and further development. Nevertheless, since both H13 and RB5 are biological entities by nature, it is possible they can get oxidized and hence careful considerations for proper storage and handling is required to expect the desired biological activity. However, considering the urgent need for developing EGFRvIII-specific targeting molecules to tackle refractory tumors, we believe that our work provides a reliable platform for further research and development in the area of EGFRvIII-targeted peptide therapeutics. From a translational viewpoint, synthesizing EGFRvIII-targeting small peptides in sterile conditions may pose challenges compared to developing small molecule inhibitors. However, we believe these challenges can be overcome by synthesizing the small peptides in bulk quantities, leveraging advancements in peptide manufacturing technology.

## 5. CONCLUSION

To summarize, we have developed novel active-targeting peptides that can specifically recognize and bind to tumorigenic EGFRvIII receptors of refractory cancers such as glioblastoma. Among the peptides identified using RPL and mAb, H13 and RB5 peptides showed appreciable recognition and binding towards both purified EGFRvIII ECD and EGFRvIII-positive glioma. Notably, compared to H13, RB5 showed lesser binding towards wild-type EGFR. Further, although both H13 and RB5 neither affected the cell viability nor interfered the phosphorylation of EGFRvIII and PI3K in glioma cells, RB5 has better possibility for efficient tumor tissue penetration due to its relatively small size. These evidences indicate that RB5 is a better choice compared to H13 peptide and may be utilized as EGFRvIII-targeting agent. Importantly, RB5 has strong potential for theranostic applications in refractory cancers, if cross-linked with NIR/PDT dyes (near-infrared photothermal/photodynamic therapy) [31], MRI/CT contrast agents (for real-time, non-invasive early cancer diagnosis and chemotherapy monitoring), nuclear/radiolabeling (radioactive/RFA therapy) or nuclear imaging moieties (in PET imaging). Finally, RB5 holds strong potential to selectively eliminate EGFRvIII expressing cancers, if covalently conjugated with antitumor agents/therapeutic nanoparticles [32].

## Author contributions

Sunitha K V: Conceptualization, Methodology, Formal analysis, Investigation, Writing-Original draft preparation, Visualization; Giridharan L M: Conceptualization, Methodology, Formal analysis, Investigation, Writing-Original draft preparation, Visualization; Krishnakumar Menon: Conceptualization, Formal analysis, Writing - Review & Editing, Supervision, Funding acquisition; Lakshmi Sumitra V: Conceptualization, Formal analysis, Writing - Review & Editing, Supervision, Funding acquisition.

## Acknowledgments

We sincerely thank Amrita Centre for Nanosciences and Molecular Medicine, Kochi, India for providing the infrastructural support for the successful accomplishment of this research work.

## Conflicts of interest

The authors declare no conflicts of interest.

## Funding

This work was supported by Department of Biotechnology (DBT), Government of India (Grant No. 6242-P2lIRGCBIPMD/DBT/LSVC/2015). S.K.V. and G.L.M. received senior research fellowships (SRF) individually from Council of Scientific and Industrial Research (CSIR), Govt. of India.

## Notes

### Competing Interest Statement

The authors have declared no competing interest.

