## Supplementary information for "Outsmarting refractory glioblastoma using *in silico* guided, phage display derived peptides targeting oncogenic EGFRvIII"

**Lakshmi Sumitra Vijayachandran***

School of Nanosciences and Molecular Medicine, Amrita Institute of Medical Sciences, Amrita Vishwa Vidyapeetham, Kochi, 682 041, India.

### 1. Materials and Methods

#### 1.1. Materials

##### 1.1.1. Chemicals and Reagents

All chemicals were purchased from Sigma Aldrich (MO, USA) or Merck (NJ, USA), unless otherwise specified. Protease inhibitor cocktail was purchased from Sigma Aldrich (MO, USA). Protein marker and Coomassie stain were purchased from Bio-Rad (CA, USA). Anti-His antibody was purchased from Abcam (MA, USA) and EGFRvIII antibody was procured from Bioss (USA). Ni-NTA resin was procured from Qiagen (Hilden, Germany). Chemiluminescence substrate and PVDF membrane were purchased from Millipore (MA, USA). Anti-M13 Antibody and Histrap Crude FF column were obtained from GE healthcare (IL, USA). DMEM (Dulbecco's Modified Eagle Medium) and penicillin-streptomycin were purchased from Thermo scientific (MA, USA). Antibiotics for bacterial culture, IPTG (Isopropyl β-D-1-thiogalactopyranoside), LB (Luria Bertani) media, and 2XYT media were procured from Himedia Laboratories (Maharashtra, India). FBS (Fetal Bovine Serum) was bought from Panbiotech (Aidenbach, Germany). Immunocytochemistry (ICC) kit was procured from Biogenex (CA, USA). MTS reagent was procured from Promega (WI, USA). EGFR and PI3K antibodies were purchased from Cell signaling (MA, USA). Anti-rabbit IgG antibody was bought from Santa Cruz Biotechnology (TX, USA). Free peptides and biotinylated peptides were synthesized and procured from Genscript (NJ, USA) as per the novel amino acid sequences that we obtained via sequencing. Unless otherwise mentioned, all chemicals and reagents were utilized as received.

##### 1.1.2. Bacterial strains, Plasmid vectors, and Phage display system

*E. coli* ER2738 was a kind gift from Prof. Martin Reick (Amrita School of Biotechnology, Kerala, India). MultiBac system that was developed in European Molecular Biology Laboratory (Grenoble, France) was a kind gift from Dr. Imre Berger. M13K07 and Ph.D.™ -12 random peptide phage display library were procured from New England Biolabs (MA, USA). *E. coli* Top10F' strain was procured from Invitrogen (CA, USA) and pCDisplay4 vector was purchased from Creative Bioscience (NY, USA).

**1.1.3. Cell lines**

U87-MG (EGFR-positive glioma cells), C6 (EGFR-negative glioma), and CTX TNA2 (normal astrocytes) were procured from ATCC (VA, USA). U87-MG.ΔEGFR (EGFR-positive and EGFRvIII-positive) cell line was a kind gift from Dr. Terrance Johns, Monash University (Australia). All mammalian cell lines were cultured under sterile conditions in DMEM supplemented with 10% FBS, 100 Unit/mL penicillin, and 100 mg/ml streptomycin in plasma pretreated culture dishes or T-flasks, and incubated at 37°C in 5% humidified $CO_2$ atmosphere. Appropriate culture media replenishments were provided as required, and the cells were maintained at ~80% confluence in log phase and utilized for cell studies.

**1.2. Methods**

To identify EGFRvIII binding peptides, we utilized two approaches: 1. using monoclonal antibody (mAb) template, 2. using random peptide phage display library. In the first approach, peptides based on mAb 528 sequence were expressed on phage coat protein, which were thereafter screened to identify EGFRvIII binding peptides. In the second approach, a commercially available random peptide library that was composed of 12 amino acid peptides displayed on M13 coat protein was screened to identify EGFRvIII binding peptides.

**1.2.1. Design and generation of mAb 528 based peptide displaying phages**

Six peptide sequences based on mAb 528 Fab (PDB ID: 2Z4Q) were selected to identify EGFRvIII binding peptides (Supplementary Table 1). Genes coding for these peptides were synthesized and cloned into pCDisplay4 vector in Genscript (NJ, USA). Peptide displaying phages were generated as described [1]. Briefly, each phagemid was transformed into ER2738 cells after which, the cells containing phagemid were grown in 5 ml 2XYT containing 100 µg/ml carbenicillin and 2% glucose at 30°C, 200 rpm till $O.D_{600}$=0.5. Helper phage, M13KO7 was added to each tubes and incubated at 37°C, 200 rpm, for 1 h. 2.5 ml of the culture was transferred into 25 ml

2XYT media containing 100 µg/ml carbenicillin and 50 µg/ml kanamycin and grown overnight at 30°C, 200 rpm. Culture supernatant was collected by centrifuging the culture at 6000 rpm for 15 min at 4°C. 5 ml of 20% PEG/2.5 M NaCl was added to the supernatant and incubated for 3 h. After centrifugation at 12,000 x g for 15 min at 4°C, the pellet obtained was resuspended in 1 ml Tris-buffered saline (TBS), reprecipitated with 200 µl 20% PEG/2.5 M NaCl and incubated overnight on ice. After centrifugation at 13,500 rpm for 10 min, the pellet obtained was resuspended in 200 µl TBS and incubated at 65°C for 20 min. The phages obtained were used for whole cell phage ELISA.

**1.2.2. Whole cell Phage ELISA of mAb 528 based peptides on glioma cells**

For whole cell phage ELISA, human glioma cells expressing wild-type EGFR (U87-MG) and EGFRvIII (U87-MG.ΔEGFR) were seeded (25000 cells/well) and incubated overnight for cell attachment. Thereafter, the cells were fixed with 1% paraformaldehyde (PFA) for 20 min at 4°C and blocked with 200 µl of 2% MPBS (skimmed milk in PBS) for 30 min at room temperature (~25°C). 100 µl ($10^{12}$ virions) of individual recombinant phages were added to the wells in triplicate for both the cell lines and incubated at room temperature for 2 h. After aspirating the phages, cells were washed three times with PBST (PBS/0.1% Tween- 20). 100 µl HRP conjugated anti-M13 antibody (1:5000 dilution in 2% MPBS) was added and incubated for 1 h at room temperature. After aspirating the antibody, wells were washed thrice with PBST and once with PBS. 100 µl of TMB substrate was added and incubated for 10 min at room temperature. Thereafter, optical absorbance was measured at 450 nm using a microplate reader, and average O.D. of the triplicates was plotted with standard deviation.

**1.2.3. Small scale induction and purification of H13 peptide**

pCDisplay4-H13 phagemid was transformed into *E. coli* Top10F' cells and grown in LB media supplemented with carbenicillin (100 µg/ml).To induce protein expression, IPTG (1 mM) was added to bacterial culture when the $OD_{600}$=0.5. The cultures were incubated at different temperatures and time points (30°C for 3 h, 5 h, and 12 h; 37°C for 3 h, 5 h, and 12 h) for optimization of expression. Cells were lysed by sonication and soluble fraction was run on 10% SDS PAGE gel to study H13 peptide expression. For western blot, samples were run on 10% SDS PAGE gel, blotted onto a PVDF membrane, blocked with 5% BSA (bovine serum albumin) in TBST (Tris buffered saline with 0.1% Tween-20). After probing with HRP tagged anti-His antibody (1:10000 dilution), blot was washed with TBST (3 x 10 min).The bands were detected by adding chemiluminescence substrate and imaged in Bio-Rad Chemidoc imaging station (CA, USA).

For small scale purification of H13, 50 ml bacterial culture was induced with 1 mM IPTG for 3 h at 37°C. Purification was performed using Qiagen Ni-NTA resin under hybrid conditions. Briefly, cells were lysed in 1 ml of lysis buffer (100 mM $NaH_2PO_4$, 10 mM Tris Cl, 8 M urea, pH 8.0) and the clarified cell lysate was incubated with the resin for 3 h. After collecting the flow through, resin was washed twice with 500 µl denaturing wash buffer (100 mM $NaH_2PO_4$, 10 mM Tris·Cl, 8 M urea pH 6.3.), and then with 500 µl native wash buffer (50 mM $NaH_2PO_4$, 300 mM NaCl, 20 mM imidazole, pH 8.0). Elution was carried out three times in 100 µl native elution buffer A (50 mM $NaH_2PO_4$, 300 mM NaCl, 250 mM imidazole, pH 8.0) followed by two elutions with 500 µl native elution buffer B (50 mM $NaH_2PO_4$ , 300 mM NaCl, 500 mM imidazole, pH 8.0). All the fractions were analyzed on 10% SDS PAGE and western blot using anti-His antibody.

#### 1.2.4. Purification of H13 peptide

To scale up the purification, clarified cell lysate from a 250 ml culture was gradually diluted to 25 ml with lysis buffer. Cell lysate was filtered using 0.22 µm syringe filter and applied to HisTrap CRUDE FF column connected to the AKTA START FPLC system, using pump at a flow rate of 5 ml/min. Thereafter, the column was washed with 10 column volume (CV), denaturing wash buffer, followed by 3 CV native wash buffer. Elution was carried out with 3 CV elution buffer A and 2 CV elution buffer B. The elute was collected in eight 1 ml fractions followed by 15 ml fractions and analyzed using SDS PAGE and western blot. To further enrich the protein, elute obtained from Ni affinity purification was run on a 10% SDS PAGE gel. After staining with coomassie blue, the bands corresponding to H13 peptide were cut using a scalpel blade and pooled into a single tube. Gel was digested in cold potassium phosphate buffer (pH 7.2) and probe sonicated till the gel liquefied completely [2]. The tube was centrifuged at 13,500 rpm for 30 min at 4°C. The gel contents were thereafter transferred to a 3 kDa cut-off column and centrifuged at 13,500 rpm for 10 min. After washing once with 100 µl potassium phosphate buffer, 100 µl of PBS was added to the filter and the peptide was collected by centrifugation at 13,500 rpm for 2 min.

**1.2.5. Biopanning, titration, amplification, and ssDNA isolation**

We have utilized the Ph.D.$^{TM}$-12 phage display peptide library that is based on a combinatorial library of random dodecapeptides, fused to the coat protein pIII of M13 bacteriophage. The 12-mer peptides are displayed at the N-terminus of pIII and a short spacer (Gly-Gly-Gly-Ser) connects the coat protein to the peptide. The library is composed of ~$10^9$ electroporated sequences amplified once to yield ~100 copies of each sequence [3].

To identify EGFRvIII binding peptides, two methods of biopanning were employed: a) subtractive biopanning and b) reverse biopanning. In subtractive biopanning, each round of biopanning was composed of initial panning on wild-type U87-MG cells, followed by biopanning on U87-MG.ΔEGFR cells. For this, $10^{12}$ phages suspended in DMEM were added to U87-MG cells grown to confluency in a 12-well plate and incubated for 1 h in the cell culture incubator. The supernatant that contained the unbound phage was added to the confluent monolayer of U87-MG.ΔEGFR cells in a 12-well plate and incubated for 2 h. Cells were washed once with wash buffer (1 ml $PBS^+$ with 0.1% BSA). After the washing step, the cells were lysed by adding a hypotonic buffer (1 ml of 30 mM Tris–HCl, pH 8.0) followed by one freeze thaw cycle (-80°C for 20 min followed by thawing at room temperature) . 10 µl of this eluate was used for titration, and the remaining phages were amplified and used for the next round of biopanning. Stringency in each round was improved by increasing Tween-20 concentration in the wash buffer from 0.1% to 0.5%. In the final step, the cell surface bound phages (acid fraction) were collected using an acid wash buffer (0.1 M HCl-Glycine, pH 2.2 + 0.9% NaCl). Cells were lysed by adding the hypotonic buffer, and one freeze thaw cycle in -80°C to collect the output fraction.

In reverse biopanning, phages resuspended in DMEM media were added to U87-MG.ΔEGFR cells grown to confluency in a 12-well plate. After 1 h of incubation, the cells were lysed to collect the phages using the hypotonic buffer, as mentioned above. These phages were amplified and then used for panning against U87-MG cells. The unbound fraction was used for biopanning against U87-MG.ΔEGFR cells. Washing and elution of phages from U87-MG.ΔEGFR cells were done using the same procedure for subtractive biopanning. In the third round, the phages that were unbound to U87-MG cells were collected and amplified. Titration was carried out after each round of biopanning using manufacturer's protocol and plaque forming unit/10 µl was calculated. Single plaques obtained from the third round of both subtractive and reverse biopanning were amplified using manufacturer's protocol. Briefly, the plate that has countable number of plaques was selected to pick individual phages. From the subtractive biopanning output, 38 plaques; acid fraction, 20 plaques; and reverse biopanning output, 35 phages were obtained. 500 µl of overnight culture of ER2738 was added to 20 ml of 2XYT+Tetracycline (20 µg/ml). Phage plaques to be amplified were added to the 20 ml ER2738 culture and incubated at 200 rpm for 4.5 h at 37°C. The culture was transferred to an oak ridge tube and centrifuged for 10 min at 12000 x g at 4°C. The phages were isolated using PEG precipitation method and incubated at 65°C for 20 min to kill bacterial cells. 20 phages from the acid fraction, 34 phages from the output fraction of subtractive biopanning, and 32 phages from the reverse biopanning output were selected for whole cell ELISA on U87-MG.ΔEGFR cell line to further screen for phages with higher affinity for EGFRvIII. Amplified phages were used in whole cell phage ELISA on U87-MG.ΔEGFR cells according to the protocol mentioned in 2.2.2. ssDNA was isolated from phages selected based on whole cell phage ELISA results in order to identify the peptide displayed by these.

For ssDNA isolation, each phage was amplified and the final phage pellet was resuspended in 100 µl phenol-chloroform, vigorously mixed, and centrifuged at 13,500 rpm for 10 min. The supernatant was collected in a fresh centrifuge tube, precipitated using 90% ethanol and 3 M sodium acetate, and incubated at -20°C for 2 h. The pellet obtained was washed with 70% ethanol and resuspended in 30 µl Tris buffer (pH 8.0). 5 µl of the ssDNA was analyzed on 1% agarose gel. 20 µl of the ssDNA was submitted for sequence analysis in Scigenom Labs, Kochi, India. This ssDNA sequence was compared with the M13 genome sequence available in the Ph.D.™ phage display library to get the peptide encoding DNA sequence. The peptide sequence was obtained using the Translate tool (ExPASy) [4].

#### 1.2.6. *In silico* molecular modeling and docking studies

The tertiary structure of EGFRvIII ectodomain (EGFRvIII ECD) was homology modeled using Modeler [5] employing wild-type EGFR (PDB ID: 1IVO) as the template, and ligand-binding sites of EGFRvIII were identified using Q-SiteFinder [6]. For the modeling of peptides (SB12, SB31, RB5), a *de novo* approach involving hidden Markov model-derived structural alphabet was used [7,8]. Briefly, from the amino acid sequence of peptides, structural alphabet profiles that could describe the conformations of four consecutive residues were predicted, which was thereafter assembled using a greedy algorithm and employing a modified optimized potential for efficient protein structure prediction coarse-grained force field. Following that, starting from an amino acid sequence, a series of simulations were performed to obtain the most representative, energetically favorable conformations. Using signatures of 25 peptides having 9-23 amino acids, the structures of peptides with lowest energy conformations differing by 2.6 A˚, were generated. Structural validation of the modeled protein and peptides were studied using Ramachandran plot [9]. Peptide-EGFRvIII ECD binding was studied employing peptide-protein docking simulations and the interactions within 4 Aº distance between the ligands and receptor were analyzed.

#### 1.2.7. *In vitro* studies

##### 1.2.7.1. Peptide ELISA on cell lines

U87-MG, U87-MG.ΔEGFR, or CTX TNA2 cells were seeded (25000 cells/well) in 96-well plates and following the cell attachment (overnight incubation), the cells were gently washed with PBS, fixed using 1% PFA and incubated at 4°C for 20 min. The fixed cells were washed thrice with PBS, blocked using 1% BSA (in PBS) and incubated for 30 min at room temperature. Biotinylated peptides were added to the cells at predetermined concentrations and incubated for 2 h at room temperature. Following that, the wells were washed thrice with PBST (PBS containing 0.05% Tween-20) and 100 µl of 1 µg/ml Streptavidin-HRP conjugate was added and incubated for 90 min. After washing the wells four times with TBST, 100 µl TMB was added and incubated for 20 min. Finally, after adding 100 µl 1N HCl (reaction termination), the optical absorbance at 450 nm was recorded using a microplate reader. Average O.D. of the triplicates was plotted with standard deviation.

##### 1.2.7.2. Cytotoxicity of anti-EGFRvIII peptides

The cytotoxicity of anti-EGFRvIII peptides (SB12, SB31, RB5, and H13) was studied using MTS assay. Briefly, U87-MG.ΔEGFR or CTX TNA2 cells were seeded at a density of 5000 cells/well in a 96-well plate. Following the cell attachment, the existing media was replenished with fresh media containing varying concentrations of SB12, SB31, or RB5 peptides (0.1, 1, 10, 20, and 50 µM) or purified H13 peptide (0.5, 1, 2, and 5 µg/ml) and incubated for 24 h. Untreated U87-MG.ΔEGFR or CTX TNA2 cells in culture media served as negative control. After 24 h of incubation, 120 µl MTS reagent was added to all the wells and incubated further for 4 h. Following that, optical density (490 nm) of all the cells was quantified using a microplate reader (BioTek, PowerWave XS, USA).

#### 1.2.7.3. Immunocytochemistry

For immunocytochemistry, U87-MG or U87-MG.ΔEGFR or CTX TNA2 cells were seeded in a 12-well plate to attain 30-40% confluency after overnight incubation. The cells were washed twice with PBS, fixed using 2% PFA (4°C, 20 min incubation), and washed again with PBS thrice. Blocking was performed using Biogenex kit reagents (Power block and peroxidase block, 15 min incubation with each). Following that, biotinylated peptide was added to the cells and incubated for 30 min. Next, streptavidin-HRP was added to the cells and incubated for 30 min. For staining, DAB chromogen was added to the cells and incubated for 10 min. For nuclear staining, one drop of hematoxylin was added to the cells, incubated for 2 min, and thereafter washed thoroughly using tap water. The stained cells were then visualized using an optical microscope.

#### 1.2.7.4. Peptide binding to EGFRvIII: effect on phosphorylation

To study the effect of H13 and mAb 528 on the phosphorylation of downstream targets of EGFR/EGFRvIII or U87-MG.ΔEGFR cells were cultured to attain confluency in a 12-well plate and treated with 300 nM of the antibody or peptide for 1 h. For the preparation of cell lysate, the cells were washed twice with ice cold PBS and 200 µl of whole cell lysis buffer (10 mM Tris base, 250 mM NaCl, 30 mM Na pyrophosphate, 50 mM Na fluoride, 100 µM Na orthovanadate, 2 mM iodoacetic acid, 5 µM $ZnCl_2$, 1 mM PMSF, 0.5% triton-X-100, 10% glycerol, protease inhibitor cocktail) was added to each well. Thereafter the cells were scraped out from the plate and collected into a microcentrifuge tube after one freeze (-80°C for 20 min) thaw cycle and lysed using a probe sonicator (amplitude: 37 A for 5 s). After incubation on ice for 20 min, microcentrifuge tubes containing the cells were centrifuged at 13,500 rpm for 5 min at 4°C. Following that, cell lysates (supernatant) were collected and run on 10% SDS PAGE gel. Western blot was performed using antibodies against EGFR, pEGFR, PI3K, and pPI3K. α-tubulin served as the loading control.

#### 1.2.7.5. ELISA on recombinant EGFRvIII ECD

Purified EGFRvIII ECD or negative control antigen (BSA) was coated on the surface of Nunc Immunosorb plates (10 mg/ml) and incubated overnight. After washing once with PBST, H13 peptide (50 nM, 100 nM, 200 nM, 500 nM) or RB5 peptide (10 nM, 100 nM, 1 µM, 10 µM) was added to the wells (in triplicates) and incubated for 30 min. After washing twice with PBST, 100 µl HRP conjugated anti-His antibody (1:1000) was added to H13 peptide treated wells, and 100 µl of 1 µg/ml Streptavidin HRP was added to the RB5 peptide treated wells, and incubated for 1 h. After washing the wells thrice with PBST, 100 µl TMB was added and incubated for 20 min. 100 µl 1N HCl served to terminate the reaction. Finally, absorbance at 450 nm was recorded using a microplate reader.

#### 1.2.7.6 Whole cell ELISA to compare binding affinity of mAb 528 and H13

H13 peptide and its parent antibody mAb 528 were biotinylated using ThermoEZ link-PEG-NHS-biotin, according to the manufacturer's protocol. For whole cell ELISA, U87-MG or U87-MG.ΔEGFR cells were seeded at 25,000 cells per well and fixed using PFA. After the blocking step, 100 µl of predetermined concentrations of biotinylated antibody or peptide were added to the cells and incubated for 1 h at room temperature (~26ºC). After the washing step, 100 µl of 1 µg/ml streptavidin-HRP was added and incubated for 1 h at room temperature. Following that, 100 µl of TMB substrate and 1N HCl were added and the optical absorbance was measured at 450 nm using a plate reader. Average O.D. of the triplicates was thereafter plotted with standard deviation.

| Sl No. | Peptide | Peptide sequence |
|---|---|---|
| 1 | L12 | DILMTQTPLSLPVSLGDQASISCRSSQNIVHNNGITYLEWYLQRPGQSPKLLIYKVSDRF |
| 2 | L13 | DILMTQTPLSLPVSLGDQASISCRSSQNIVHNNGITYLEWYLQRPGQSPKLLIYKVSDRFSGVPDRFS GSGSGTDFTLKISRVEAEDLGI |
| 3 | L14 | DILMTQTPLSLPVSLGDQASISCRSSQNIVHNNGITYLEWYLQRPGQSPKLLIYKVSDRFSGVPDRFS GSGSGTDFTLKISRVEAEDLGIYYCFQGSHIPPTFGGGTKLEIKRADAAPTV |
| 4 | H23 | SYWMHWVKQRHGHGPEWIGNIYPGSGGTNYAEKFKNKVTLTVDRSSRTVYMHLSRLTSED |
| 5 | H13 | QVQLQQSGSEMARPGASVKLPCKASGDTFTSYWMHWVKQRHGHGPEWIGNIYPGSGGTNYA EKFKNKVTLTVDRSSRTVYMHLSRLTSED |
| 6 | H14 | H1QVQLQQSGSEMARPGASVKLPCKASGDTFTSYWMHWVKQRHGHGPEWIGNIYPGSGGTN YAEKFKNKVTLTVDRSSRTVYMHLSRLTSEDSAVYYCTRSGGPYFFDYWGQGTSLTVSSAK |

Supplementary Table 1: Sequence of mAb 528 based peptides

| **Element Binding property** | **EGFR vIII interacting residues** |
|---|---|
| Cationic binding (metals/non-metals) | 16C, 30D, 36K, 38C, 41P, 169D, 233V, 235C, 265I, 267C, 289I, 291C, 235Y |
| Metal ion binding | 16C, D30, 36 K, C38, P41, D169, V233, C235, I265, C267, I289, C291, Y235, C300, A306, Y335 |
| Ion binding (cationic/anionic) | C16, E26, D30, K36, C38, P41, I49, D169, P227, V233, C235, N261, I265, C267, I289, C291, C300, A306, 327H, 329C, 334T, 335Y |
| Transition metal ion binding (Fe, Cu, Zn, Ni, etc) | C16, D30, K36, C38, P41, H127, P227, V233, C235, R256, E263, I265, C267, I289, C291, C300, P305, V325, H327,L328, C329 |

Supplementary Table 2: Amino acid residues involved in metal binding activity of EGFRvIII.

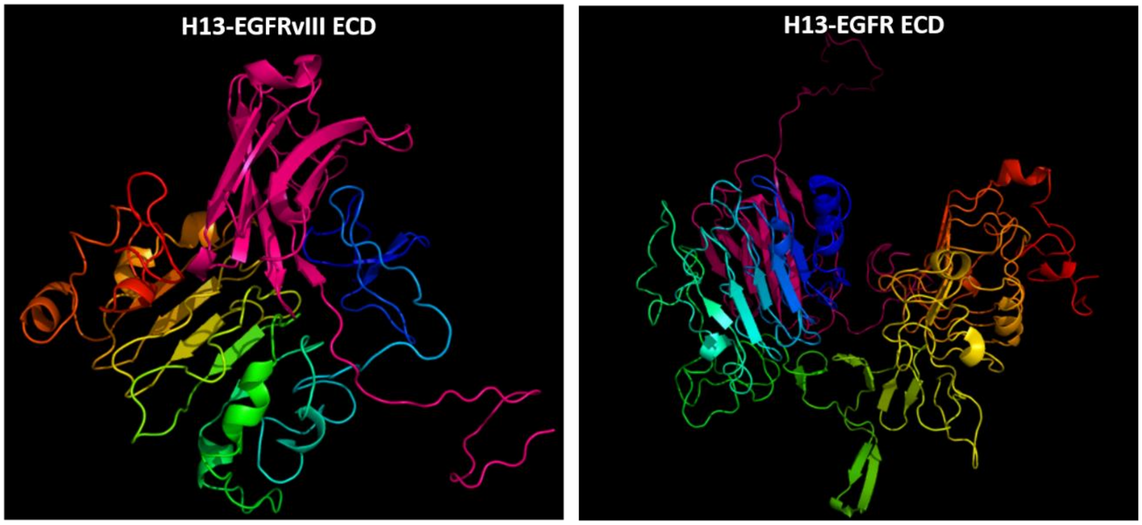


| Peptide-protein docked complex | Binding energy (kcal/mol) |
|---|---|
| H13-EGFRvIII ECD | -16.9 |
| H13-EGFR ECD | -12.6 |

Supplementary figure 1: *In silico* docking simulations of H13 with EGFRvIII and EGFR ECD. Binding energies are indicated in the table.

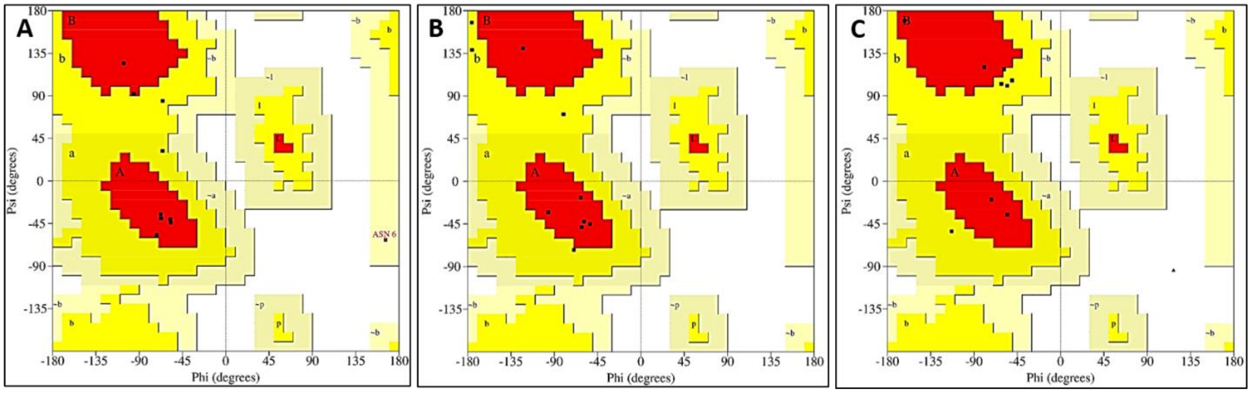


Supplementary Figure 2: Structural verification of A) SB12, B) SB31, and C) RB5 phages using Ramachandran plot.

| EGFRvIII residues | Residue number | SB12 | Residue number |
|---|---|---|---|
| GLN | 145 | VAL | 253 |
| LEU | 62 | ASN | 255 |
| ASN | 179 | MET | 260 |
| GLN | 121 | LYS | 256 |
| SER | 177 | THR | 252 |
| VAL | 154 | ILE | 251 |
| SER | 155 | LEU | 257 |
| LYS | 202 | ALA | 250 |
| GLN | 121 | LEU | 257 |
| HIS | 146 | VAL | 253 |
| VAL | 154 | THR | 252 |
| SER | 177 | ALA | 250 |
| ILE | 204 | ILE | 251 |
| GLN | 121 | ASN | 255 |
| PRO | 86 | PRO | 258 |
| HIS | 146 | THR | 252 |
| PRO | 86 | LYS | 256 |
| HIS | 83 | PRO | 254 |
| ILE | 204 | ALA | 250 |
| VAL | 87 | LYS | 256 |
| LEU | 119 | ASN | 255 |
| GLN | 145 | PRO | 254 |
| LEU | 119 | VAL | 253 |
| LEU | 85 | LYS | 256 |
| GLY | 178 | ILE | 251 |
| GLN | 145 | THR | 252 |
| ILE | 175 | ILE | 251 |
| SER | 155 | MET | 260 |
| ILE | 175 | THR | 252 |
| PHE | 149 | THR | 252 |
| GLN | 121 | VAL | 253 |
| LEU | 85 | ASN | 255 |
| SER | 155 | PRO | 258 |
| ILE | 175 | ALA | 250 |
| ALA | 152 | VAL | 253 |
| SER | 177 | ILE | 251 |
| PRO | 86 | LEU | 257 |
| GLN | 121 | PRO | 254 |
| GLN | 121 | PRO | 258 |
| SER | 205 | HIS | 261 |
| GLY | 178 | MET | 260 |
| LYS | 180 | MET | 260 |
| HIS | 83 | ASN | 255 |
| VAL | 154 | LEU | 257 |
| GLN | 148 | THR | 252 |
| ALA | 152 | THR | 252 |
| LEU | 119 | PRO | 254 |
| VAL | 154 | VAL | 253 |

| EGFRvIII residues | Residue number | SB31 | Residue number |
|---|---|---|---|
| GLY | 35 | LYS | 261 |
| GLU | 113 | THR | 258 |
| LYS | 112 | ASP | 250 |
| ARG | 47 | PRO | 257 |
| ASN | 135 | ASP | 253 |
| THR | 110 | ASP | 253 |
| GLU | 32 | ARG | 260 |
| ILE | 138 | ARG | 260 |
| GLU | 137 | TYR | 251 |
| SER | 170 | ARG | 260 |
| VAL | 36 | ARG | 260 |
| ARG | 246 | ASP | 250 |
| GLU | 168 | ARG | 260 |
| LYS | 109 | HIS | 252 |
| GLU | 32 | LEU | 259 |
| GLU | 134 | ASP | 250 |
| ILE | 138 | LEU | 259 |
| ASN | 135 | ASP | 250 |
| ARG | 164 | TYR | 251 |
| GLU | 30 | THR | 258 |
| GLY | 35 | ARG | 260 |
| GLY | 195 | LYS | 261 |
| SER | 165 | TYR | 251 |
| THR | 76 | LEU | 256 |
| MET | 31 | THR | 258 |
| LYS | 112 | SER | 255 |
| LEU | 166 | TYR | 251 |
| ARG | 47 | SER | 255 |
| THR | 76 | PRO | 257 |
| ASN | 11 | ARG | 260 |
| SER | 165 | ASP | 250 |
| GLU | 137 | LEU | 259 |
| GLU | 32 | LYS | 261 |
| ASN | 74 | ASP | 253 |
| ILE | 138 | LYS | 261 |
| ARG | 37 | ARG | 260 |
| GLU | 137 | LEU | 256 |
| LYS | 167 | LYS | 261 |
| GLU | 32 | THR | 258 |
| ILE | 138 | THR | 258 |
| GLU | 30 | PRO | 257 |
| SER | 77 | THR | 258 |
| ASN | 135 | HIS | 252 |
| GLU | 168 | LYS | 261 |
| MET | 31 | PRO | 257 |
| LYS | 112 | TYR | 251 |
| TYR | 29 | PRO | 257 |
| GLU | 113 | LEU | 259 |

| EGFRvIII residues | Residue number | RB5 | Residue number |
|---|---|---|---|
| GLY | 162 | ALA | 251 |
| THR | 215 | LEU | 261 |
| ARG | 246 | LEU | 254 |
| GLY | 162 | ARG | 257 |
| ASP | 129 | ARG | 257 |
| ASN | 135 | ALA | 251 |
| GLY | 216 | LEU | 261 |
| ARG | 234 | PRO | 255 |
| GLU | 134 | THR | 250 |
| SER | 160 | PRO | 260 |
| HIS | 131 | TYR | 253 |
| PRO | 231 | PRO | 260 |
| ASP | 235 | ARG | 257 |
| THR | 159 | PRO | 260 |
| CYS | 183 | LEU | 261 |
| CYS | 212 | LEU | 261 |
| SER | 160 | LEU | 261 |
| GLU | 134 | ALA | 251 |
| TRP | 229 | ARG | 257 |
| GLU | 232 | PRO | 260 |
| THR | 128 | ARG | 257 |
| PRO | 231 | LEU | 261 |
| THR | 159 | LEU | 261 |
| ARG | 164 | LYS | 252 |
| GLU | 134 | LYS | 252 |
| GLU | 247 | PRO | 255 |
| GLY | 162 | LYS | 252 |
| SER | 165 | ALA | 251 |
| THR | 128 | PRO | 260 |
| GLN | 217 | LEU | 261 |
| HIS | 131 | THR | 250 |
| LYS | 109 | TYR | 253 |
| ASP | 235 | ALA | 251 |
| ARG | 234 | TYR | 253 |
| ASP | 129 | PRO | 260 |
| ARG | 234 | ALA | 251 |
| THR | 128 | GLY | 259 |
| ALA | 132 | TYR | 253 |
| THR | 128 | LYS | 252 |
| LEU | 161 | ARG | 257 |
| HIS | 131 | LYS | 252 |
| ASP | 129 | LYS | 252 |
| LYS | 109 | THR | 250 |
| SER | 160 | ARG | 257 |
| LEU | 130 | LYS | 252 |
| ARG | 234 | THR | 250 |
| PRO | 231 | ARG | 257 |
| GLU | 232 | ARG | 257 |
| ARG | 234 | LEU | 254 |

Supplementary Figure 3: Closely interacting residues of SB12, SB31, and RB5 peptides with EGFRvIII ECD.
